# Archaeogenetics of Late Iron Age Çemialo Sırtı, Batman: Investigating maternal genetic continuity in North Mesopotamia since the Neolithic

**DOI:** 10.1101/172890

**Authors:** Reyhan Yaka, Ayşegül Birand, Yasemin Yılmaz, Ceren Caner, Sinan Can Açan, Sidar Gündüzalp, Poorya Parvizi, Aslı Erim Özdoğan, Zehra İnci Togan, Mehmet Somel

## Abstract

North Mesopotamia has witnessed dramatic political and social change since the Bronze Age, but the impact of these events on its demographic history is little understood. Here we study this question by analysing the recently excavated Late Iron Age settlement of Çemialo Sırtı in Batman, southeast Turkey. Archaeological and/or radiocarbon evidence indicate that the site was inhabited during two main periods: the first half of the 2nd millennium BCE and the first millennium BCE. Çemialo Sırtı reveals nomadic items of the Early Iron Age, as well as items associated with the Late Achaemenid and subsequent Hellenistic Periods. Mitochondrial DNA (mtDNA) haplotypes from 12 Çemialo Sırtı individuals reveal high genetic diversity in this population, conspicuously higher than early Holocene west Eurasian populations, which supports the notion of increasing population admixture in west Eurasia through the Holocene. Still, in its mtDNA composition, Çemialo Sırtı shows highest affinity to Neolithic north Syria and Neolithic Anatolia among ancient populations studied, and to modern-day southwest Asian populations. Population genetic simulations do not reject continuity between Neolithic and Iron Age, nor between Iron Age and present-day populations of the region. Despite the region’s complex political history and indication for
increased genetic diversity over time, we find no evidence for sharp shifts in north Mesopotamian maternal genetic composition within the last 10,000 years.

## 1. Introduction

North Mesopotamia became densely populated around 12,000 BCE, after the Last Glacial Period (Rosenberg and Erim-Özdoğan, 2011). Both during the Pre-pottery Neolithic A (PPNA) and Pre-pottery Neolithic B (PPNB) the region was a favourable geographical zone for sedentary and nomadic hunter-gatherers. Besides excavated PPNA sites (*e.g.* Tell Aswad, Tell Qaramel, Djade El-Mughara, Mureybet, Jerf el Ahmar, Göbekli Tepe, Hallan Çemi, Hasankeyf Höyük, Çayönü, Qermez Dere, Nemrik) many regional settlements have been detected in surveys. Most of these PPNA sites were abandoned by the PPNB (except for a few exceptions, *e.g.* Tell Aswad, Mureybet, Tell Qaramel, and Çayönü), while many new, large and complex sedentary communities emerged during the PPNB across north Mesopotamia and the Levant, between ca.7600-6200 BCE (*e.g.* Tell Ramad, Tell Halula, Bouqras, Tell ain el-Kerkh, Abu Hureyra, Tell Seker al-Aheimar, Tell Fekheriye, Cafer Höyük, Nevali Çori, Mezraa Teleilat). The beginning of crop and animal domestication, earliest metallurgy, various burial practices, increasing in usage of Anatolian obsidian, special buildings with particular items and decorations were fundamental features of the PPNB period. By ca.6600-6200 cal. BCE, however, with the 8.2 ka cold/arid event, rapid climate change-driven social and economic effects led to the decline of PPNB societies in the Levant, north Mesopotamia and Anatolia, resulting either in abandonment of settlements or their reduction in size (Staubwasser and Weiss 2006). Archaeological data indicate that the Neolitization of the Balkans and the Caucasus also occurred during this time period, between ca.6600-6000 cal. BCE (Weninger et al 2006; Weninger et al 2009; Weninger et al 2014).

The exact timing of the beginning of the Pottery Neolithic (PN) in north Mesopotamia is still debated. In addition to the early excavated PN sites in Syria and Iraq (*e.g.* Tell Şeker al Aheimar, Jarmo, Kül Tepe, Tell Hassuna), recently dug Sumaki Höyük in the north of Lower Garzan Basin and Salat Camii Yanı at Bismil-Diyarbakır also display changes in land choice from PPNB (Erim-Özdoğan 2011). After temperature and precipitation conditions improved, north Mesopotamia was densely reoccupied, with the Halaf culture (cal. 5400-4990 BCE), carrying both sedentary and semi-nomadic characters, flourishing in the region. The Halaf was eventually replaced by the Ubeid culture (5500-4000 BCE) of south Mesopotamian origin, followed by another south Mesopotamian originated culture, Uruk. The Uruk controlled the region until ca.3200 BCE, when the 5.2 ka event caused their collapse through severe drought (Staubwasser and Weiss 2006). Subsequently, small settlements of local Late Chalcolithic cultures spread across the region up to the Caucasus (Marro 2010). Between 2500-1500 BCE, the Holocene 4.2 ky cold/arid event was likely the main factor in the collapse of Akkadian Empire (ca.2350-2150 BCE). Following the 4.2 ka event, the abandoned lands of north Mesopotamia and Palestine were reoccupied intensively and reorganized by sedentarized nomadic pastoralists (Staubwasser and Weiss 2006). Meanwhile, the region witnessed major political changes (Nissen and Heine 2003): Neo-Assyrian influence (10th-7th century BCE), was followed by that of Babylon (6th century BCE), the Achaemenid Empire of Persia (6th-4th century BCE), and the empires of Alexander and the Seleucid of Hellen origin (4th-2nd century BCE). Between 2nd century BCE and 7th century CE, the region was dominated by Rome/Byzantium in the west, and Parthians and later Sasanians in the east. North Mesopotamian region underwent Muslim Arab conquests in the 8th century, and from the 11th century onward, it fell under the rules of central and east Asian-related groups, including the Seljuks, Mongols, and the Ottomans. Notably, these major political changes have also been attributed to more recent rapid climate change and demographic effects (Büntgen et al., 2016; Kuzucuoğlu, 2015)

Still, north Mesopotamia’s demographic history remains little known. One major question is whether rapid climate change-driven population fluctuations, as during the end of the PPNB, involved population replacement from distant regions. Such replacement would appear as allele frequency shifts in the regional gene pool. Another question involves whether political and cultural change was accompanied by major gene flow events, or whether immigration was a weak force relative to social forces. The genetic legacy of migration and resettlement policies of various imperial states (including Assyrians, Seleucids, Romans, and Ottomans; see Schechla, 1993; Zürcher 2009; Nissen and Heine 2003; Woolf 2015), is likewise unclear.

Ancient DNA (aDNA) today holds great potential to address these questions, and methods have improved to the point that aDNA from temperate regions can be recovered with reasonable success. Yet, there has been only a single study of north Mesopotamia archaeogenetics published to date. This work investigated human mitochondrial DNA (mtDNA) from PPNB sites from north Syria, Tell Halula and Tell Ramad (Fernández et al., 2014). Ten individuals’ haplotypes were obtained, which showed high similarity (*i.e.* low *F_ST_*) to the mtDNA composition of modern-day populations of the Near East, Caucasus and south Europe, with the highest similarity to Cyprus. Still, the authors of this study noted clear differences in haplotype frequencies between the Neolithic north Syrian population and those of modern-day southwest Asia. They suggested that such shifts could arise due to immigration or genetic drift, although they did not address these hypotheses explicitly.

Here we present archaeological and genetic data from Çemialo Sırtı, Batman, Turkey, a culturally diverse Late Iron Age site in north Mesopotamia. Comparing Çemialo Sırtı mitochondrial DNA sequences with those of ancient and modern-day west Eurasian populations we study admixture and continuity in the demographic history of north Mesopotamia.

## 2. Archaeology of Çemialo Sırtı

Çemialo Sırtı (CML) is located in the territory of the Yazıhan Village, Gedikli hamlet of the Beşiri district in Batman Province, southeast Turkey (Fig. 1, SI Fig. 1). The site is located in the middle of the Lower Garzan Basin watered by the Garzan Stream, one of the main tributaries of the Tigris River. The site lies on a low ridge approximately 10-12 m above the plain at an elevation of 503-512 m on the south bank of the Garzan Stream, at the junction of Malabini and the Garzan Streams. It is within the direct impact area of the Ilısu Dam, which started to be constructed in the early 2000’s. Rescue excavations were started in 2009 and continued between 2013-2015 at two sites, Gre Amer and Çemialo Sırtı. An area of 3600 m^2^ has been exposed at the site displaying material culture elements characteristic of two periods: the first half of the 2nd millennium BCE and the 1st millennium BCE.

**Fig. 1.**
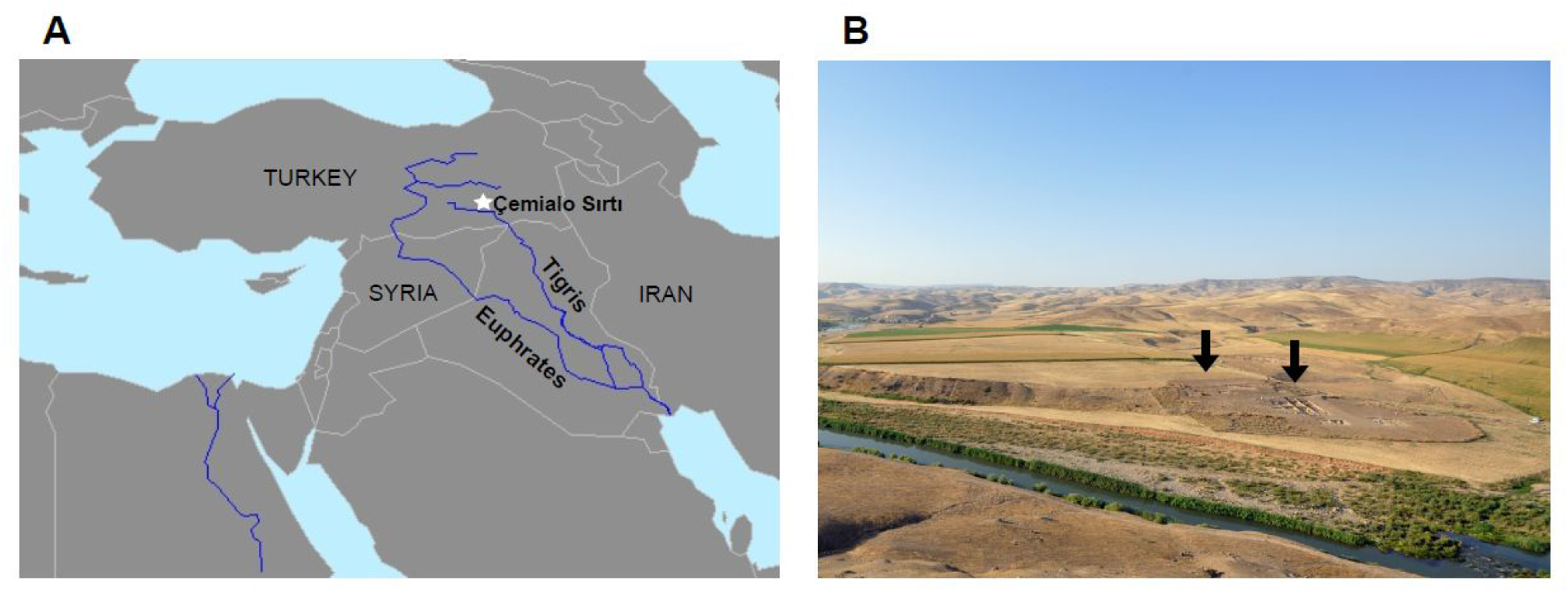
The geographic location **(A)** and aerial view **(B)** of Çemialo Sırtı, Batman. Black arrows show the different regions of the excavation site.

During the Artuqid Period (12th-14th centuries CE), the Lower Garzan Plain appears to have played an active role in the commercial route system between Mesopotamia, Iran, and Caucasus, as well as west and north Anatolia (Supplementary Information). Roman and Byzantine presence in the region is less evident, but according to surface surveys (Erim-Özdoğan and Sarıaltun 2011) and results from rescue excavations at Gre Amer and Çemialo Sırtı, the Garzan Plain was densely occupied during the 1st millennium BCE. Both Gre Amer and Çemialo Sırtı display material culture evidence from diverse traditions, detected not only in architecture but also in artifacts of daily use and in burial practices.

In Çemialo Sırtı, the Late Achaemenid Period (550–330 BCE) (Sampson, 2008; Curtis and Tallis, 2005; Ahmad et al., 2013) presence is identified by Triangle Ware, as well as typical pottery forms and bronze objects like kohl sticks with castellated heads known from the Achaemenid layers of the sites Nimrud, Deve Hüyük, Kamid el-Loz, Tell Jigan, and Pasargadae (Curtis, 2004). Çemialo Sırtı also yields Hellenistic footprints, represented by a silver *Tetra Drahmi* of Alexander the Great (ca. 325-323 BCE), pottery sherds, a head of a terracotta figurine. Traces of temporal architectural elements, pits, and items such as dog burials and crude pottery, generally attributed to Early Iron Age nomadic peoples, have also been discovered (Supplementary Information).

A total of 47 graves have been excavated at Çemialo Sırtı, including pithos/jar burials (*n*=19), stone cists (*n*=10) and pit graves (*n*=18) (SI Figs. 2-3). Most individuals were placed contracted on their left or right sides, with or without burial goods. Burial characteristics include the use of stone cists and pit graves common in north Mesopotamia since the Bronze Age, such as reported for Titriş Höyük and Lidar Höyük (Algaze et al., 1995; Hauptmann, 1982, 1984, 1987, 1993; Matney and Algaze, 1995), and these features seem to have continued into the Late Achaemenid Period. One exception is an individual with ornaments suggesting Early Iron Age cultures, such as stone bell-shaped hollowed amulet carried by a copper wire. Different burial practices have been recorded in different sites of the eastern and western parts of the Near East during the Hellenistic period, such as flexed inhumation in Nimrud, Iraq (Oates and Oates, 1957), and cremation, flexed, or extended inhumation in Antandros, west Turkey (Yağız, 2009); some of the graves at CML, such as SK35, a body laid on its back holding an imitation of a black glazed plate, can also be attributed to Hellenistic Period. Further information on the site and the graves are provided in the Supplementary Information and SI Figs. 1-3.

## 3. Materials and Methods

### 3.1. Samples

The samples reported in this study were recovered during excavations at Çemialo Sırtı site in 2013-2014 field season in accordance with aDNA study guidelines. We analyzed 16 individuals excavated from three different locations: conglomerates on top of the ridge, around the middle of hillside, and on the skirt of the ridge (Supplementary Table S1). All were primary burials and dated to the Late Iron Age based on archaeological context, except one individual whose ornaments implied an earlier period (see section 2). We used 22 bone and tooth samples from 16 human skeletons for DNA extractions (eight female adults, three male adults, three pre-adults, and two adults whose sex was undetermined).

Three of these skeletons were radiocarbon dated to 535-395 cal. BCE, 395-205 cal. BCE, and 390-205 cal. BCE (95% probability) by Beta Analytic, Miami, USA (Supplementary Table S1). Thus, all three potentially overlap with the Late Achaemenid Period (550–330 BCE), and the latter two may also overlap with the subsequent Hellenistic Period (330-31 BCE).

### 3.2. Contamination avoidance

Genetic analyses were carried out at the aDNA facilities of the Middle East Technical University (METU). We used all the precautions to avoid contamination (Gilbert et al., 2005; Olivieri et al., 2010; Ottoni et al., 2011; Paabo et al., 2004): using samples cleanly stored at site without washing, using a dedicated aDNA laboratory in a building physically separated from the post-PCR laboratory and under strictly controlled conditions, following irreversible steps throughout aDNA extraction, PCR and post-PCR laboratory procedures, using negative controls at each step of the experimental processes, independent repeats of experiments at multiple times for reproducibility. Genetic profiles of team members (all archaeologist, anthropologist, and laboratory researchers involved) were also determined for comparison with genetic profiles of ancient individuals to avoid contamination from modern human DNA. Details on contamination avoidance procedures are provided in the Supplementary Information.

### 3.3. Ancient DNA extraction, amplification, and authenticity assessment

After decontamination and pulverization of tooth and/or phalanx samples, 0.2-0.3 g of powder was digested in a proteinase-K lysis buffer and DNA extracted through silica-based spin columns following Ottoni et al. (2011). The first and second hypervariable regions (HVRI-HVRII) of mtDNA were amplified and sequenced in multiple independent experiments using five and two overlapping fragments (ranging from 109–166 bp in size) to read 357 bp, between nucleotide positions (np) 16009-16365, and 217 bp, between np 49–265, respectively, for the HVI and HVII regions (Ottoni et al., 2011).

PCR amplification procedures and cycling conditions followed Ottoni et al. (2011) with slight modifications. We conducted minimum three independent extractions from each sample, and three independent amplifications from each extraction. We required clear sequences that were consistent across all the overlapping fragments from independent extract and PCR repeats. Samples from four individuals, for which these attempts produced less than 3 consistent sequences, were discarded. Sequences were confirmed by considering reads from both strands of the mtDNA and determined from overlapping HVRI and HVRII fragments. We confirmed that none of the mtDNA haplotypes of research team members matched the ancient haplotypes (Supplementary Table S8). We further recorded the frequency and types of nucleotide changes among repeated sequence reads for each individual, especially C->T transitions, which is the hallmark of postmortem damage in authentic aDNA (Sawyer et al., 2012; Olivieri et al., 2010; Paabo et al., 2004). Details regarding sample preparation, aDNA extraction and amplification reactions, the list of primers used, purification and sequencing processes, precautions taken to minimize contamination and criteria adopted for assessing authenticity are provided in the Supplementary Information and Supplementary Table S7.

### 3.4. mtDNA haplogroup determination and population genetic analysis

mtDNA haplogroups of successfully amplified Çemialo Sırtı individuals were determined based on the SNPs at informative nucleotide positions of HVI and HVII region sequences (Supplementary Table S2) using the Cambridge Reference Sequence (CRS) (Anderson et al., 1981) as reference sequence and mtDNA Phylotree build 17 (van Oven and Kayser, 2009).

Çemialo Sırtı haplogroups and sequences were compared with published data from 15 ancient Eurasian populations (Mesolithic Eurasia, Neolithic, Bronze Age, and Iron Age west Eurasia, as well as Roman and Byzantium) and 13 modern-day populations from west Asia and southeast Europe. These datasets were chosen to contain a minimum of 10 individuals each (Supplementary Table S3).

Multiple sequence alignments were performed using ClustalW software (Thompson et al., 1994; http://www.ebi.ac.uk/Tools/msa/). Within mtDNA HVRI, a 240 bp region was shared among all populations (between np 16126-16365 on the CRS). To measure population genetic differentiation among populations we calculated Slatkin’s linearized *F_ST_* between pairs of populations (Slatkin, 1995) using Arlequin v.3.5.2 (Excoffier and Lischer, 2010). Significance of *F_ST_* values was calculated using Arlequin by 10,000 random permutations of population labels. To visualize genetic differentiation between the populations, the pairwise *F_ST_* matrix was summarized using two-dimensional non-metric multidimensional scaling (MDS) using the “metaMDS” function in the R package “vegan” (Oksanen, 2008) and plotted in R v.3.2.4 (http://www.r-project.org/). Pairwise *F_ST_* with Çemialo for all populations was displayed on a geographical map using the “rworldmap”, “maps” and “SDMTools” packages implemented in R.

Haplotype (gene) diversity, the probability that two randomly chosen haplotypes are different in the sample (Nei and Kumar, 2000), was estimated for all population samples using Arlequin. Absolute and relative frequencies of mtDNA haplogroups were further calculated using a Bayesian calculator (http://www.causascientia.org/math_stat/ProportionCI.html) presenting the estimated frequencies together with the 95% Bayesian credible intervals (CI) in Çemialo Sırtı, Neolithic north Syria, modern-day Syria and southeastern Anatolia populations. Details about statistical analysis and populations data are given in the Supplementary Table S3 and Supplementary Information.

### 3.5. Coalescent simulations of population continuity

For specific pairs of diachronic population comparisons, a recent population *X* and a more ancient population *Y*, we tested the null hypothesis that the observed *F_ST_* value could arise solely by genetic drift (*i.e.* population continuity). We compared the observed *F_ST_* values with those generated by coalescent simulations.

For each pairwise comparison *X* and *Y*, separated by a certain number of generations, we simulated DNA sequence data using the fastsimcoal2 simulation software (Excoffier et al., 2013). We used a demographic model of exponential growth (*e.g.* Ottoni et al., 2016; Sverrisdóttir et al., 2014) employing a range of parameter values, *i.e.*, different effective population sizes (Supplementary Table S5). The exponential growth rate was calculated as the natural logarithm of the ratio of the population sizes of *X* and *Y*. We assumed a generation time of 25 years, and the mutation rate of 3.6x10^-6^ per site per generation for the human mtDNA HVRI (Richards et al., 2000). We performed 1000 simulations for each combination of population sizes, summing up to ~250,000-400,000 simulation runs for each comparison (see section 3.5, Fig. 4A). In each simulation, we sampled from the population serially through time under the exponential population growth scenario (Sverrisdóttir et al., 2014).

We carried out 1000 simulations for each combination of population sizes, calculated *F_ST_* using Arlequin v.3.5.2 (Excoffier and Lischer, 2010), and calculated the proportion of simulated *F_ST_* values that are greater than the observed *F_ST_*. If the observed *F_ST_* was found within the range of *F_ST_* values from the simulations, this would be compatible with population continuity between the two populations (including migration from genetically similar populations), or could indicate a lack of statistical power. Conversely, if most simulations yielded *F_ST_* values smaller than that observed, the null hypothesis of continuity can be rejected (assuming that the demographic scenario and parameters are valid). Such rejection could be interpreted as gene flow and/or migration from genetically distinct third populations that led to differentiation of *X* from *Y*, or equivalently, that *X* from *Y* did not belong to the same ancestral population.

## 4. Results

We studied a total of 16 Çemialo Sırtı individuals’ remains (Supplementary Table S1). Of these, three randomly chosen individuals were radiocarbon dated to between 535-205 cal. BCE, consistent with archaeological evidence for Late Achaemenid (550–330 BCE) (Sampson, 2008; Curtis and Tallis, 2005; Ahmad et al., 2013) presence in Çemialo Sırtı (section 2). In the rest of this study we therefore assume that these individuals represent a Late Iron Age north Mesopotamian population.

We obtained reproducible mtDNA HVRI-HVRII (HVI and HVII regions) sequences from 12 out of 16 individuals’ remains, including 9 teeth and 3 phalanx samples (75% success rate). Authenticity was assessed by requiring at least 3 overlapping fragments from independent extract and PCR repeats, by the presence of cytosine deamination indicative of postmortem decay, and lack of match to mtDNA haplotypes of the research team (section 3.3). All ancient mitochondrial DNA sequences produced in this study have been deposited in the NCBI GenBank database (accession number pending).

### 4.1. mtDNA haplogroup composition of Çemialo Sırtı

We determined mtDNA haplogroups for the 12 Çemialo Sırtı individuals (Table 1 and Supplementary Table S2). The most common haplogroup was H (*n*=5), one of the most abundant types in present-day Europe and in the Near East (Achilli et al., 2004; Richards et al., 2000). We also observed subtypes of haplogroup R (R2 and R6, *n*=1 each), M (M1a1, *n*=1), and HV (HV and HV8, *n*=1 each). One individual belonged to subtype of N1(N1a1a1a), a common haplogroup in Neolithic farmer populations (Brandt et al., 2014; Szécsényi-Nagy et al., 2015), and another belonged to a haplogroup common in European Paleolithic and Mesolithic populations, U2 (Bramanti et al., 2009; Brandt et al., 2014; Der Sarkissian et al., 2013; Krause et al., 2010). Çemialo Sırtı haplogroup frequency credible intervals largely overlapped with those from Neolithic north Syria (*n*=10), modern-day southeast Anatolia (n=224), and modern-day Syria (*n*=48) (Table 1). An exception to this pattern was macrohaplogroup M, which was absent in both the Neolithic and modern-day Syrian datasets. We found no obvious association between haplogroup type and burial characteristics (pithos, location, or presence of burial gift) (Supplementary Table S1).

**Table 1.** Haplogroup frequencies (N) together with the 95% Bayesian credible intervals (CI) estimated for Late Iron Age Çemialo Sırtı, Neolithic north (N) Syria, and for modern-day southeast (SE) Anatolia and Syria population samples. Data sources are described in Table S3 .

|  | Late Iron Age<br>Çemialo Sirtı |  | Neolithic<br>N Syria |  | Modern-day<br>SE Anatolia |  | Modern-day<br>Syria |  |
| --- | --- | --- | --- | --- | --- | --- | --- | --- |
| Haplogroup | %(N) | CI | %(N) | CI | %(N) | CI | %(N) | CI |
| A | 0 | 0-25 | 0 | 0-28 | 0.9(2) | 0-3 | 0 | 0-7 |
| B | 0 | 0-25 | 0 | 0-28 | 1.3(3) | 0-4 | 2.1(1) | 0-11 |
| C | 0 | 0-25 | 0 | 0-28 | 3.6(8) | 2-7 | 0 | 0-7 |
| D | 0 | 0-25 | 0 | 0-28 | 0.9(2) | 0-3 | 0 | 0-7 |
| G | 0 | 0-25 | 0 | 0-28 | 0.9(2) | 0-3 | 0 | 0-7 |
| H | 41.7(5) | 19-68 | 10(1) | 2-41 | 21(47) | 16-27 | 18.8(9) | 10-32 |
| HV | 16.7(2) | 5-45 | 20(2) | 6-52 | 4.5(10) | 2-8 | 8.3(4) | 3-20 |
| I | 0 | 0-25 | 0 | 0-28 | 1.8(4) | 1-4 | 0 | 0-7 |
| J | 0 | 0-25 | 0 | 0-28 | 6.7(15) | 4-11 | 6.3(3) | 2-17 |
| K | 0 | 0-25 | 40(4) | 17-69 | 5.8(13) | 3-10 | 8.3(4) | 3-20 |
| L0 | 0 | 0-25 | 0 | 0-28 | 0 | 0-2 | 0 | 0-7 |
| L | 0 | 0-25 | 10(1) | 2-41 | 0.9(2) | 0-3 | 0 | 0-7 |
| M | 8.3(1) | 2-36 | 0 | 0-28 | 0.4(1) | 0-2 | 0 | 0-7 |
| N1 | 8.3(1) | 2-36 | 0 | 0-28 | 3.6(8) | 2-7 | 2.1(1) | 0-11 |
| N* | 0 | 0-25 | 10(1) | 2-41 | 0.4(1) | 0-2 | 0 | 0-7 |
| R0 | 0 | 0-25 | 10(1) | 2-41 | 8.9(20) | 6-13 | 2.1(1) | 0-11 |
| R | 16.7(2) | 5-45 | 0 | 0-28 | 2.2(5) | 1-5 | 6.3(3) | 2-17 |
| T | 0 | 0-25 | 0 | 0-28 | 10.3(23) | 7-15 | 12.5(6) | 6-25 |
| **U1** | 0 | 0-25 | 0 | 0-28 | 1.3(3) | 0-4 | 2.1(1) | 0-11 |
| **U2** | 8.3(1) | 2-36 | 0 | 0-28 | 3.6(8) | 2-7 | 2.1(1) | 0-11 |
| **U3** | 0 | 0-25 | 0 | 0-28 | 7.6(17) | 5-12 | 12.5(6) | 6-25 |
| **U5** | 0 | 0-25 | 0 | 0-28 | 2.7(6) | 1-6 | 4.2(2) | 1-14 |
| **U7** | 0 | 0-25 | 0 | 0-28 | 4(9) | 2-7 | 2.1(1) | 0-11 |
| **U\*** | 0 | 0-25 | 0 | 0-28 | 0 | 0-2 | 6.3(3) | 2-17 |
| **W** | 0 | 0-25 | 0 | 0-28 | 5.4(12) | 3-9 | 2.1(1) | 0-11 |
| **X** | 0 | 0-25 | 0 | 0-28 | 1.3(3) | 0-4 | 2.1(1) | 0-11 |
| **Total (N)** | **12** |  | **10** |  | **224** |  | **48** |  |

### 4.2. Population comparisons of genetic diversity

We compared the Late Iron Age Çemialo Sırtı population with mtDNA datasets representing 14 ancient and 13 modern-day populations from west Eurasia, chosen to include at least 10 individuals (Fig. 2 and Supplementary Table S3). The 14 published ancient mtDNA datasets comprised populations from Mesolithic west Eurasia (Bramanti et al., 2009; Der Sarkissian et al., 2013), Pre-Pottery Neolithic northern Syria (Tell Halula and Tell Ramad) (Fernández et al., 2014), Pottery Neolithic northwest Anatolia (Barcın) (Mathieson et al., 2015), Early Neolithic Central Europe (LBK, Starčevo, LBKT) (Brandt et al., 2014; Szécsényi-Nagy et al., 2015), Early and Late Neolithic southwest Europe (northeast Iberia and Spain Treilles culture) (Gamba et al., 2012; Lacan et al., 2011), Bronze Age Crete (Minoan) (Hughey et al., 2013), Early Bronze Age Central Europe (Unetice culture) (Brandt et al., 2014), as well as Roman and Byzantine Period southwest Anatolia (Sagalassos) (Ottoni et al., 2011, 2016). This ancient dataset contained 467 individuals in total, including Çemialo Sırtı individuals (*n*=10-88 individuals per population, median=24). The modern-day populations were chosen from west Asia and south Europe (Alkan et al., 2014; Brisighelli et al., 2012; Comas et al., 1996; Comas et al., 2004; González et al., 2007; Irwin et al., 2008; Nasidze and Stoneking, 2001; Ottoni et al., 2011, 2016; Roostalu et al., 2007; Schönberg et al., 2011; Serin et al., 2016; Vernesi et al., 2001), summing up to a total of 1483 individuals (*n*=30-286 individuals per population, median=90).

**Fig. 2.**
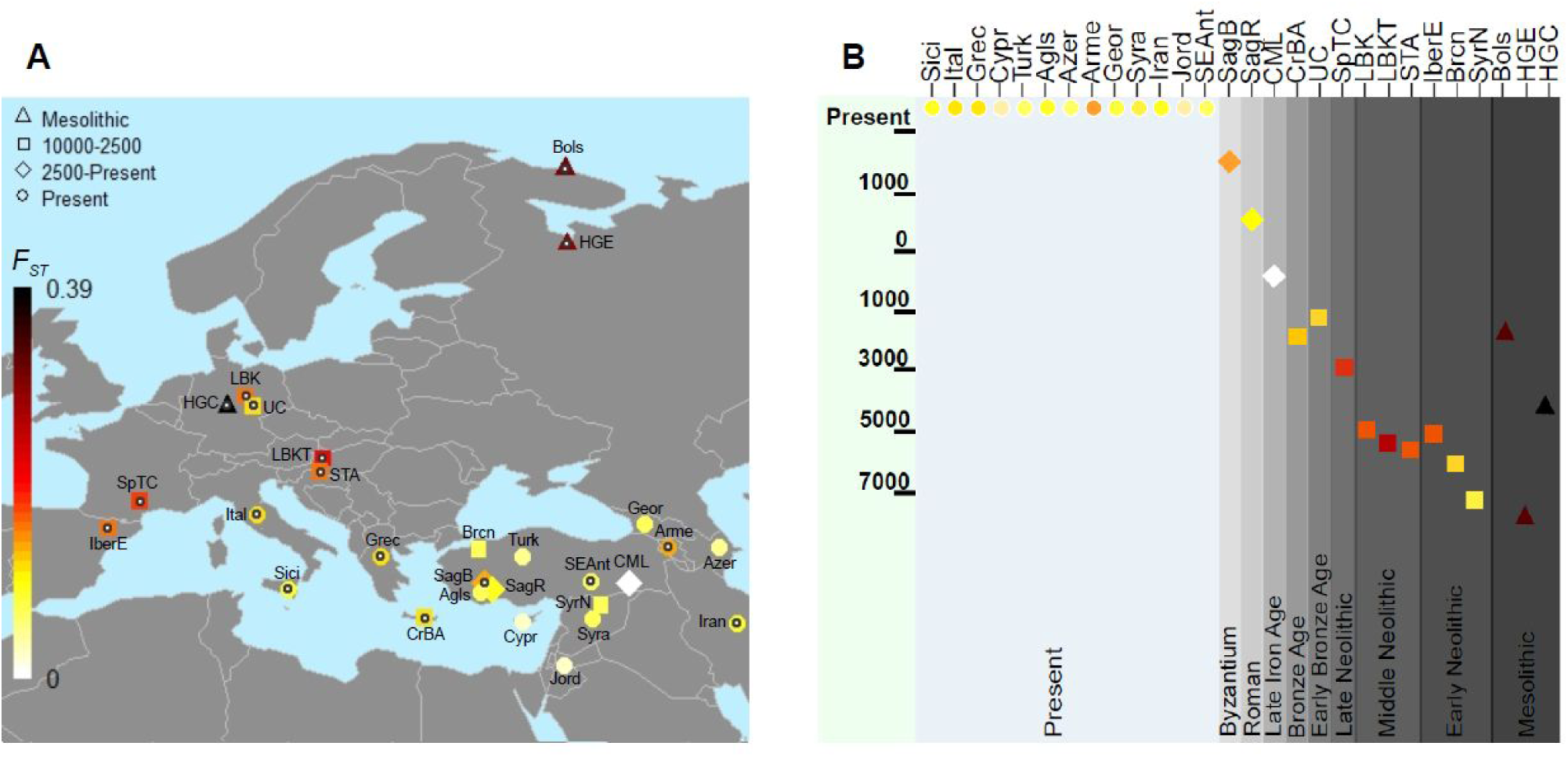
Mitochondrial DNA differentiation (*F_ST_*) between the Çemialo Sırtı population and ancient or modern-day populations from west Eurasia. **(A)** The map shows the locations of the 15 ancient and 13 modern-day populations from west Eurasia. Shapes indicate the age or archaeological context of the population, the colors indicate level of *F_ST_* with Çemialo Sırtı according to the key on the left. Empty black circles indicate significant differentiation, as measured by random permutations (p < 0.05). **(B)** The approximate ages of the populations included in the study (y-axis), with colors indicating *F_ST_* with Çemialo Sırtı (see Table S3 for more detail on the populations). Modern-day populations; Agls: southwest Anatolia (Aglasun), Arme: Armenia, Azer: Azerbaijan, Ital: Italy, Grec: Greece, Cypr: Cyprus, Geor: Georgia, Iran: Iran, Jord: Jordan, SEAnt: southeast Anatolia, Sici: Sicily, Syra: Syria, Turk: Turkey. Ancient populations; Bols: Bolshoy (Mesolithic northwest Russia), Brcn: Barcın Höyük (Neolithic northwest Anatolia), CML: Çemialo Sırtı, CrBA: Crete Bronze Age (Minoan), HGC: Hunter-Gatherer of central Europe (Mesolithic central Europe), HGE: Hunter-Gatherer of east Europe (Mesolithic northwest Russia), IberE: Iberia Early Neolithic (Northeast Iberia), LBK: Linear Pottery culture (Neolithic central Europe), LBKT: Linearbandkeramik culture in Transdanubia (Neolithic central Europe), SagB: Sagalassos Byzantium (Southwest Anatolia), SagR: Sagalassos Roman (Southwest Anatolia), SpTC: Spain Treilles culture (Late Neolithic west Europe), STA: Early Neolithic Starčevo culture (Neolithic central Europe), SyrN: Neolithic north Syria, UC: Unetice culture (Early Bronze Age central Europe).

We estimated haplotype diversity within all populations based on a 240 bp HVRI region shared among all sequences (Supplementary Table S6). Diversity estimates for ancient Eurasian populations (0.818-0.989, median=0.927) were noticeably lower than those for modern-day Eurasians (0.959-0.995, median=0.977) (Mann-Whitney U-test p<0.001, assuming independence among populations). Our observation that modern-day west Eurasian populations carry higher genetic diversity than ancient west Eurasians groups agrees with earlier reports based on nucleotide diversity and runs of homozygosity estimated from ancient and modern whole genomes (Jones et al., 2015; Kılınç et al., 2016; Skoglund et al., 2014). Interestingly, Late Iron Age Çemialo Sırtı had the highest haplotype diversity estimate among the 15 ancient populations, and modern-day Syria (but not Turkey or southeast Anatolia) had the highest diversity estimate among the 13 contemporary populations. The lowest haplotype diversity levels were found for Mesolithic European groups and in Neolithic north Syria.

### 4.3. Population differentiation

We calculated pairwise *F_ST_*, an estimate of between-population genetic differentiation relative to within-population diversity, among all 15 ancient and 13 modern populations’ mtDNA sequences, and tested statistical significance by random permutations. *F_ST_* values between Late Iron Age Çemialo Sırtı and other populations are depicted in Fig. 2 and Fig. 3. *F_ST_* values between Çemialo Sırtı and other ancient Eurasian populations (range=0.026-0.389, median=0.068) were generally higher than those between Çemialo Sırtı and modern populations (range=0.009-0.056, median=0.033), which is expected given low diversity in ancient Eurasian populations (section 4.2).

**Fig. 3.**
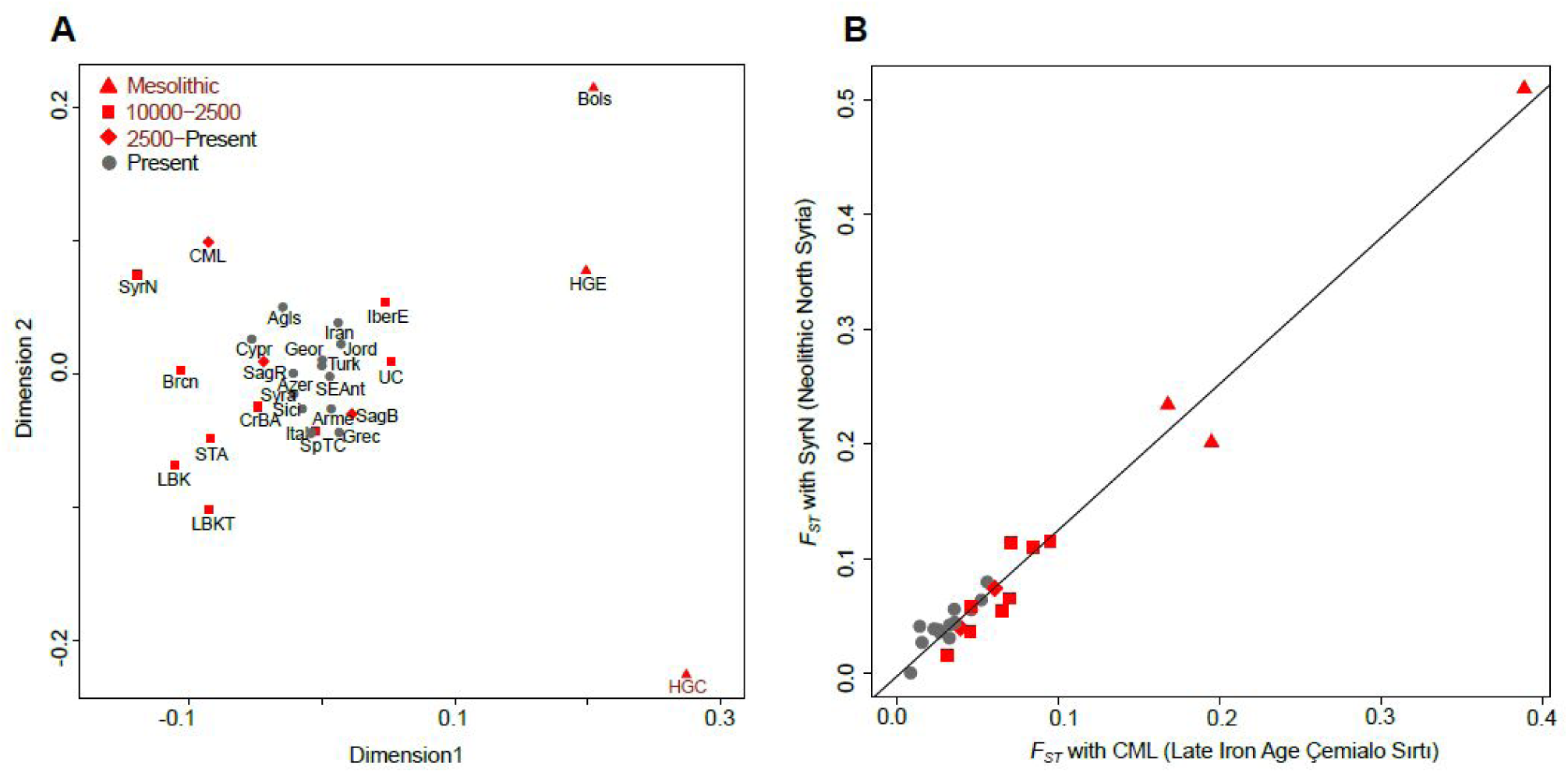
Mitochondrial DNA differentiation across ancient and modern-day populations. **(A)** Multidimensional scaling (MDS) plot summarizing mtDNA *F_ST_* measures across 15 ancient and 13 modern-day populations from west Eurasia (Table S3). The abbreviations follow Fig. 2. The MDS calculation had a stress value of 0.13. **(B)** *F_ST_* between Çemialo Sırtı (x-axis) or the north Syrian Neolithic (y-axis) with third populations.

Among ancient populations, the mtDNA composition of the Çemialo Sırtı was closest to those of Neolithic north Syria and to Neolithic northwest Anatolia, followed by Roman Period Anatolia and Bronze Age Crete. Çemialo Sırtı showed higher differentiation from Early Neolithic European Neolithic populations and to Byzantine Period Anatolian, and was most differentiated from European Mesolithic populations. Among modern-day populations, Çemialo Sırtı had lowest *F_ST_* with Cyprus, followed by Jordan, Azerbaijan, Turkey and Syria, and higher *F_ST_* values with Greece, Italy, and Armenia. Çemialo Sırtı mtDNA composition thus showed higher affinity to those of geographically proximal populations of the last 10,000 years, with increasing genetic differentiation upon increasing distance from north Mesopotamia. Byzantine Period Anatolia (Sagalassos) and modern-day Armenia were the only exceptions to this pattern, being geographically close but genetically more differentiated (Fig. 2). Among the *F_ST_*-based comparisons between Çemialo Sırtı and ancient or modern-day populations, 11/14 and 6/13 were significant as measured by random permutations, respectively (*p*<0.05, Fig. 2, Supplementary Table S4; we note that statistical significance is a function of both the level of differentiation and of sample size).

We further summarized pairwise mitochondrial *F_ST_* values among all 15 ancient and 13 modern-day populations using multidimensional scaling (Fig. 3A). As with similar summaries of ancient and modern nuclear genomic variation of west Eurasian populations (e.g. Lazaridis et al., 2014; Skoglund et al., 2014), the main coordinates set apart early/mid Holocene groups, while more recent populations were represented in the middle, implying increasing population admixture over time (Kılınç et al., 2016; Lazaridis et al., 2016). In the multidimensional scaling plot, the first dimension separated ancient populations of Near Eastern origin (including early Neolithic Central Europe) and Mesolithic European populations. The second dimension set apart Central Europe (LBK and Mesolithic Central Europe) from Mesolithic east Europe and Neolithic southwest Asia. The *F_ST_* measures summarised in the MDS plot also underscored the close affinity between the Late Iron Age Çemialo Sırtı population and that of Neolithic north Syria, both from the same region but separated by over seven millennia (Fig. 3).

### 4.4. Testing population continuity with Çemialo Sırtı

Although the above analysis hints at population continuity in the region, we found statistically significant differentiation in comparisons of Late Iron Age Çemialo Sırtı versus a number of neighboring modern-day populations: Iran, southeast Anatolia, Greece, and Armenia (*F_ST_*=0.036, 0.033, 0.046, and 0.056, respectively; permutation test *p*<0.05). If we assume that Çemialo Sırtı was part of a wide regional population comprising west Asia and southeast Europe during the Late Iron Age, then these modern-day populations must have differentiated from Çemialo Sırtı over the past ~2500 years (~100 generations). Three forces could account for such differentiation: natural selection, migration, and/or genetic drift. Selection on human mitochondrial DNA has been reported (Balloux et al., 2009) but strong selection in resident human populations within such short time frame is unlikely. By contrast, population differentiation by migration from genetically distinct populations is a plausible factor, especially given the region’s history of frequent social and political change (section 1). Finally, genetic drift is a ubiquitous force in natural populations. If it can be shown that genetic drift cannot solely explain differentiation, one may presume the role of migration, or that the two diachronic samples did not belong to the same population.

To address this, we conducted population genetic simulations using the fastsimcoal2 simulation software to test whether the observed differentiation between Çemialo Sırtı and a modern-day population can be explained purely by genetic drift within the same gene pool within a given time interval, assuming an exponential growth model, and a wide range of population sizes. We obtained no evidence to reject continuity between Çemialo Sırtı versus Syria, Iran, or southeast Anatolia (SI Fig. 4B, C, F), such that for the majority of parameter value combinations the observed *F_ST_* was within the null *F_ST_* distribution (>99% of parameter value combinations with *p*>0.10, Fig. 4). There was only partial evidence to reject continuity with Armenia (observed *F_ST_*=0.0493, 27% of parameter value combinations had *p*<0.10), and strong evidence to reject continuity with Greece (observed *F_ST_*=0.0437, 85% of parameter value combinations had *p*<0.10, SI Fig. 4G). The difference between comparisons with Armenia and with Greece appears to be due to sample size and hence statistical power differences (Supplementary Table S5).

**Fig. 4.**
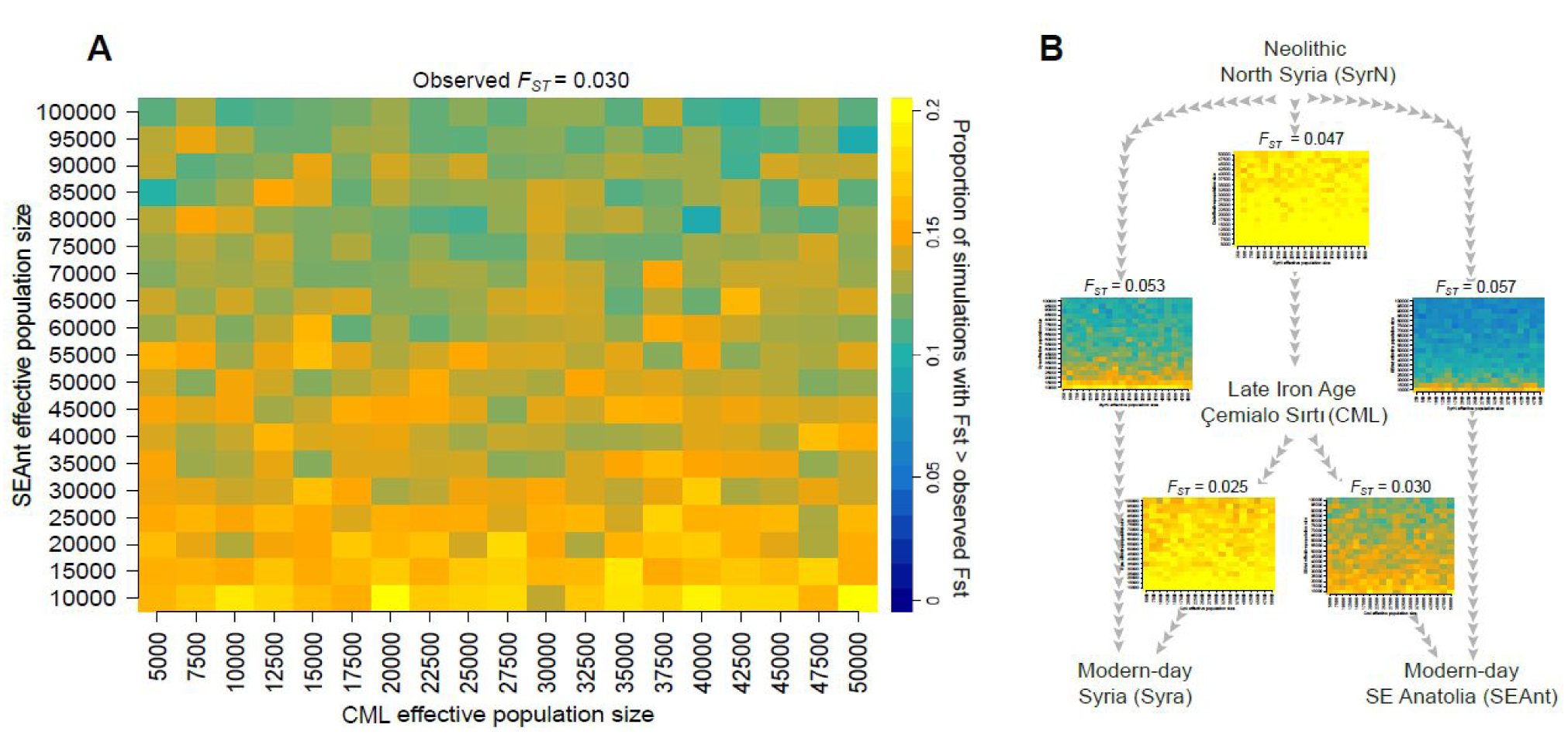
Regional population continuity tested by simulation. **(A)** The grid represents the results of 19 X 19 X 1000 serial coalescent simulations under an exponential growth model, starting with Late Iron Age Çemialo Sırtı (CML) and ending with modern-day southeast Anatolia (SEAnt). The effective population sizes of CML and SEAnt used in the simulations are shown on the x- and y-axes. The parameter ranges were chosen following (Sverrisdottir et al. 2014). The colors indicate the proportion of 1000 simulations in each grid that had *F_ST_* values greater than that observed (*F_ST_*=0.030), with warm colors indicating that the observed *F_ST_* can arise solely by genetic drift. The colder colors indicate that the observed *F_ST_* is less likely to have arisen by genetic drift, implying migration. **(B)** Results from 5 comparisons involving Neolithic north Syria, Later Iron Age Çemialo Sırtı, and modern-day Syria and SE Anatolia.

Using the same approach we next tested continuity between Çemialo Sırtı and ancient west Eurasians. Here we could not reject continuity with Neolithic north Syria and northwest Anatolia, or with Roman Anatolia (>99% of parameter value combinations with *p*>0.10). In contrast, simulations suggested a general lack of continuity in comparisons of Çemialo Sırtı versus European Mesolithic and Early Neolithic populations, as well as Byzantine Anatolia (>50% of parameter value combinations with *p*<0.10) (Supplementary Table S5). The exceptions to this pattern were Early Neolithic Iberia (observed *F_ST_*=0.071), and Early Bronze Age Central Europe (Unetice Culture, observed *F_ST_*=0.046), for which continuity with Çemialo Sırtı could not be rejected (Supplementary Table S5). Again, this is due to low sample size of the Early Neolithic Iberia dataset.

### 4.5. Testing population continuity with Neolithic north Syria

To gain better understanding into the demographic history of north Mesopotamia, we further studied the relationships between PPNB north Syria individuals (*n*=10, Fernandez et al. 2014) and other west Eurasian groups. Comparing PPNB north Syria with modern-day populations we found no significant differentiation with Cyprus, Azerbaijan, and Georgia (*F_ST_*<0.05, *p*>0.05), low but significant differentiation with Syria, Jordan, southeast Anatolia, and Sicily (*F_ST_*<0.05, *p*<0.05), and moderate differentiation with modern-day Greece, Iran, and Armenia (0.05<*F_ST_*<0.10, *p*<0.05) (Supplementary Table S5). We conducted simulations comparing Neolithic north Syria with modern-day Syria, southeast Anatolia, Iran, Greece, and Armenia: continuity was rejected in all comparisons (>75% of parameter value combinations with *p*<0.10) except for modern-day Syria (77% of parameter value combinations with *p*>0.10) (Fig. 4B, SI Fig. 4D,E).

In addition, we found that Neolithic north Syria had low differentiation with Neolithic northwest Anatolia, Çemialo Sırtı, and Roman Anatolia (*F_ST_*<0.05, non-significant in permutations). Simulations did not reject continuity in either of the three comparisons (>99% of parameter value combinations with *p*>0.10). Most other comparisons between Neolithic north Syria and ancient populations had moderate to high *F_ST_* values (*F_ST_*>0.05; Supplementary Table S5).

## 5. Discussion

Material culture evidence suggests Çemialo Sırtı was a Late Achaemenid-related settlement, although earlier nomadic and later Hellenic influences are also observed. Radiocarbon data complies with this Late Achaemenid-later Hellenistic time window (535-205 cal. BCE, including all 3 dated individuals’ 95% intervals). The mitochondrial haplotype composition of the Çemialo Sırtı sample, including macrohaplogroups H, HV, M, N, R and U2, is generally similar to both earlier and later regional populations (Table 1). Macrohaplogroup H, comprising >40% of the dataset, is still one of the most abundant types in west Eurasia (Achilli et al., 2004; Richards et al., 2000). Of note, we found one instance of macrohaplogroup M - this type was previously suggested to have been introduced in the region from Central Asia by Seljuk invasions in the eleventh century CE (Ottoni et al., 2016), while our detection of M in Iron Age CML contradicts this claim. We also observe an individual with haplogroup U2, intriguing as U haplogroups belong to a lineage associated with Palaeolithic and Mesolithic individuals of Europe (Bramanti et al., 2009; Brandt et al., 2014; Der Sarkissian et al., 2013; Krause et al., 2010) Overall, Çemialo Sırtı was not only culturally diverse, but also diverse in its maternal lineages.

Although north Mesopotamia was under Persian (Achaemenid) and Hellen political and cultural influence in the late 1st millennium (section 1), Çemialo Sırtı mtDNA composition did not show exceptional affinity to either modern-day Iran, or to Greece. Instead, Çemialo Sırtı was closest to Neolithic north Syria and northwest Anatolia. Among modern-day populations, it was closest to Cyprus, Jordan, Azerbaijan, and then to SE Anatolia and Syria; a similar pattern was found for Neolithic north Syria (Fig. 3). These observations suggest that genetic differentiation that occurred in north Mesopotamian populations over the past ten millennia, from the Neolithic to the Iron Age, and to the present, was modest (*F_ST_*<0.05). Our simulations further show that regional population continuity cannot be rejected in most comparisons (SI Fig. 4), and migration need not necessarily be invoked. We could neither reject continuity in comparisons of Neolithic north Syria versus Çemialo Sırtı, nor in comparisons of Çemialo Sırtı versus modern-day Syria or southeast Anatolia (Fig. 4, SI Fig. 4A, B), implying uninterrupted continuity over ten millennia (although lack of significance is partly attributable to low sample sizes in the ancient datasets).

The notion of regional continuity appears at odds with another observation arising both from our analysis and from earlier work: throughout the Holocene, within-population genetic diversity rose in west Eurasia, possibly via admixture among genetically distinct populations (Günther and Jakobsson, 2016; Kılınç et al., 2016; Lazaridis et al., 2016; Skoglund et al., 2014). Remarkably, haplotype diversity in Late Iron Age Çemialo Sırtı was the highest among ancient populations and on a par with those of modern-day west Eurasian populations, while Neolithic north Syria had one of the lowest diversity estimates. This difference in diversity between diachronic populations from the same region implies admixture events did take place in between the Neolithic and Late Iron Age periods.

To reconcile these two observations, *i.e.* continuity and increasing diversity, we suggest that extensive population replacement [as what happened during European Neolithization (Skoglund et al., 2014)] did not take place in north Mesopotamia within the last 10,000 years. Shifts in north Mesopotamian maternal genetic composition by immigration from genetically distinct populations must have occurred only at modest scales, undetectable at our current resolution. Alternatively, if the region’s population size was constantly large, any immigrant alleles would be rapidly diluted. Thus, despite the turbulent sociopolitics of north Mesopotamian history, the maternal lineage appears to have remained largely continuous, reminiscent of population continuity reported for the Levant (Haber et al., 2016) and for Armenia (Margaryan et al., 2017). Whether this description equally applies to nuclear genomic variation remains to be shown.

## Acknowledgements

We thank the Batman Museum and Claudio Ottoni for support, all members of the METU ancient DNA, METU CompEvo and Çemialo Sırtı teams, especially Füsun Özer, Eren Yüncü, Nihan Dilşad Dağtaş and Aliye Gündüzalp for support and helpful discussions, Düzce University Archeology Department students for help, and Düzce University for the access to the DU Biological Anthropology Laboratory. This work was funded by the Scientific and Technological Research Council of Turkey (TÜBİTAK grant no. 114Z159) and METU (BAP-07-02-2015-003).

